# Defensive freezing and its relation to approach-avoidance decision-making under threat

**DOI:** 10.1101/2021.01.29.428809

**Authors:** Felix H. Klaassen, Leslie Held, Bernd Figner, Jill X. O’Reilly, Floris Klumpers, Lycia D. de Voogd, Karin Roelofs

**Author notes:** Correspondence: Felix Klaassen, Donders Centre for Cognitive Neuroimaging, Radboud University Nijmegen, Trigon building, 6525 EN Nijmegen, The Netherlands.

## Abstract

Successful responding to acutely threatening situations requires adequate approach-avoidance decisions. However, it is unclear how threat-induced states-like freezing-related bradycardia-impact the weighing of the potential outcomes of such value-based decisions. Insight into the underlying computations is essential, not only to improve our models of decision-making but also to improve interventions for maladaptive decisions, for instance in anxiety patients and first-responders who frequently have to make decisions under acute threat. Forty-two participants made passive and active approach-avoidance decisions under threat-of-shock when confronted with mixed outcome-prospects (i.e., varying money and shock amounts). Choice behavior was best predicted by a model including individual action-tendencies and bradycardia, beyond the subjective value of the outcome. Moreover, threat-related bradycardia interacted with subjective value, depending on the action-context (i.e., passive vs. active). Specifically, in action-contexts incongruent with participants’ intrinsic action-tendencies, strong freezers showed diminished effects of subjective value on choice. These findings illustrate the relevance of testing approach-avoidance decisions in relatively ecologically valid conditions of acute and primarily reinforced threat. These mechanistic insights into approach-avoidance conflict-resolution may inspire biofeedback-related techniques to optimize decision-making under threat. Critically, the findings demonstrate the relevance of incorporating *internal* psychophysiological states and *external* action-contexts into models of approach-avoidance decision-making.

---

In the face of threat, it is extra important to weigh the value of potential outcomes of our decisions to optimally deal with the potentially harming situation. Imagine just overcoming the fear to ask for a salary raise but finding your superior in a bad mood. How do we weigh a possible detrimental outcome of a scolding against a beneficial outcome of a better income? And how do we override the initial tendency to actively back-off or passively avoid the situation? While the anticipated outcomes of the decision play a decisive role in resolving such approach-avoidance conflicts, the presence of threat triggers a psychophysiological defensive state that may inform this decision. Understanding how transient psychophysiological states affect our behavior is essential to advance models of human decision-making as well as to improve interventions for anxiety patients or first responders, who are constantly forced to make decisions under threat. Yet, existing models of decision-making do not account for defensive threat states. Here we aim to bridge this gap between affective and decision sciences. Using a novel approach-avoidance conflict task, we test the link between threat-induced psychophysiological states and value-based approach-avoidance conflict decisions and their action-implementation.

Under acute threat, activation of the autonomic nervous system triggers a defense cascade. Whereas noradrenergically-driven sympathetic upregulation is associated with increased heart rate and muscular tone enabling active fight-or-flight responses, concurrent cholinergically-driven parasympathetic dominance under proximal threat results in a net heart rate deceleration (i.e., bradycardia) and motor inhibition characteristic of a freezing response (Carrive, 1993; Nijsen et al., 2000; Roelofs, 2017; Van Der Zee, Roozendaal, Bohus, Koolhaas, & Luiten, 1997). Freezing is a universal defensive response to upcoming threat observed across many species (Bolles, 1970; Rosen, 2004). In anticipation of threat, it serves as a temporary break on the cardiac and motor system, whereby its attention and perception enhancing properties are thought to affect subsequent fight-or-flight decisions through enhanced risk assessment (Blanchard, 2017; Blanchard, Griebel, Pobbe, & Blanchard, 2011; Bradley, 2009; Hagenaars, Oitzl, & Roelofs, 2014; Mobbs, Hagan, Dalgleish, Silston, & Prévost, 2015). In rodents and other animals, threat-anticipatory freezing is generated by projections from the central nucleus of the amygdala to the midbrain periaqueductal gray (PAG), which in turn effectuates immobility via medullar projections to spinal cord motor neurons and bradycardia via the vagus nerve (Calhoon & Tye, 2015; Carrive, 1993; Silva & McNaughton, 2019; Tovote et al., 2016). Recent work in humans confirmed similar neural mechanisms underlying the human freezing response (Hashemi, Gladwin, et al., 2019; Schipper et al., 2019). Importantly, in humans, this state of immobility and bradycardia has been associated with preferred visual perception of low spatial frequency features (Lojowska, Gladwin, Hermans, & Roelofs, 2015), important for fast threat detection and facilitated by direct amygdala-visual cortex projections during freezing (Lojowska, Ling, Roelofs, & Hermans, 2018). Together with the fact that freezing is stronger when active responding is possible (Gladwin, Hashemi, van Ast, & Roelofs, 2016), and that stronger freezing is associated with faster subsequent responding (Hashemi, Gladwin, et al., 2019), this shows that freezing may play an important role in decision-making under threat. It is however unclear *how* freezing affects the decision-making process.

One possibility is that freezing affects approach-avoidance conflict decisions by acting on the subjective value computations underlying the approach-avoidance conflict. So far, however, no studies have incorporated freezing (or the associated bradycardia) in value-based models of approach-avoidance conflict decisions. Previous work using gamble-based and foraging-like paradigms has resulted in valuable formal models of how individuals resolve choice conflict under outcome uncertainty by typically contrasting monetary gains vs. losses (Bach, 2015, 2017; Charpentier, Aylward, Roiser, & Robinson, 2017; Charpentier, De Neve, Li, Roiser, & Sharot, 2016; FeldmanHall, Glimcher, Baker, & Phelps, 2016; He, Zhao, & Bhatia, 2020; Tom, Fox, Trepel, & Poldrack, 2007; Tversky & Kahneman, 1992). Given the absence of formal threat induction and concurrent psychophysiological states, such paradigms may however not generalize to real-life conflict situations that involve anxious emotional states (Delgado, Jou, & Phelps, 2011). Only few studies have incorporated threat into approach-avoidance conflicts by contrasting (monetary) rewards with receiving electric shocks (Bublatzky, Alpers, & Pittig, 2017; Kirlic, Young, & Aupperle, 2017; Park, Kahnt, Rieskamp, & Heekeren, 2011) or by contrasting threat of shock with total safety (Berns, Capra, Chappelow, Moore, & Noussair, 2008). Because these studies did not take into account the psychophysiological state of the decision-maker in their value-based decision models, it remains unclear whether such states are incorporated into the computation of the subject value of the potential outcomes that underlies the decision. If freezing feeds into such outcome value computations, this could affect the decision by biasing the conflict towards increased avoidance (Ly, Huys, Stins, Roelofs, & Cools, 2014).

A second option is that the impact of freezing on the subjective value of approach-avoidance conflict decisions depends on whether an action is required. Indeed, previous studies (Gladwin et al., 2016; Löw, Weymar, & Hamm, 2015; Wendt, Löw, Weymar, Lotze, & Hamm, 2017) found that freezing was reduced in situations where no action could be taken. However, because passive paradigms typically do not involve decision-making, the role of freezing in situations that offer the possibility for passive approach-avoidance remains unclear. Making a distinction between passive and active approach-avoidance decisions is especially relevant in light of profound individual differences in passive versus active avoidance strategies across different types of anxiety disorders (Krypotos, Effting, Kindt, & Beckers, 2015), for which an explanatory human model is largely lacking. Animal models have shown that freezing-induced action inhibition can hamper the opportunity for active avoidance (Bolles, 1970; Martinez et al., 2013; Moscarello & LeDoux, 2013; Pavlova, Rysakova, Zaichenko, & Broshevitskaya, 2020). However, more recent work in humans (Gladwin et al., 2016; Hagenaars et al., 2014; Hashemi, Zhang, et al., 2019) has shown that freezing is associated with action preparation. Therefore, how freezing affects approach-avoidance conflict decisions under threat may depend on the action context.

To test these hypotheses, we developed the Passive-active Approach-avoidance Task (PAT, **Figure 1**), in which participants are required to integrate varying monetary and shock amounts into an approach-avoidance decision in both passive and active action contexts. Action contexts were created by manipulating the movement direction of the to be approached/avoided target (i.e., either away from or towards the participant icon). We operationalized freezing in the anticipation of the decision by means of heart rate. In addition, we assessed body sway (i.e., postural freezing) to confirm bradycardia as part of the threat-anticipatory freezing response in our novel task using a stabilometric force platform (Hashemi, Gladwin, et al., 2019). First, we assessed the ability of the task to induce approach-avoidance conflict by testing effects of reward and punishment on passive and active approach-avoidance conflict decisions using mixed-effects model analyses for choice behavior and response time. Then, we applied computational modeling to investigate whether freezing-related bradycardia interacts with the subjective value of avoid versus approach choices, and/or with individuals’ context-dependent passive versus active action tendencies.

**Figure 1.**
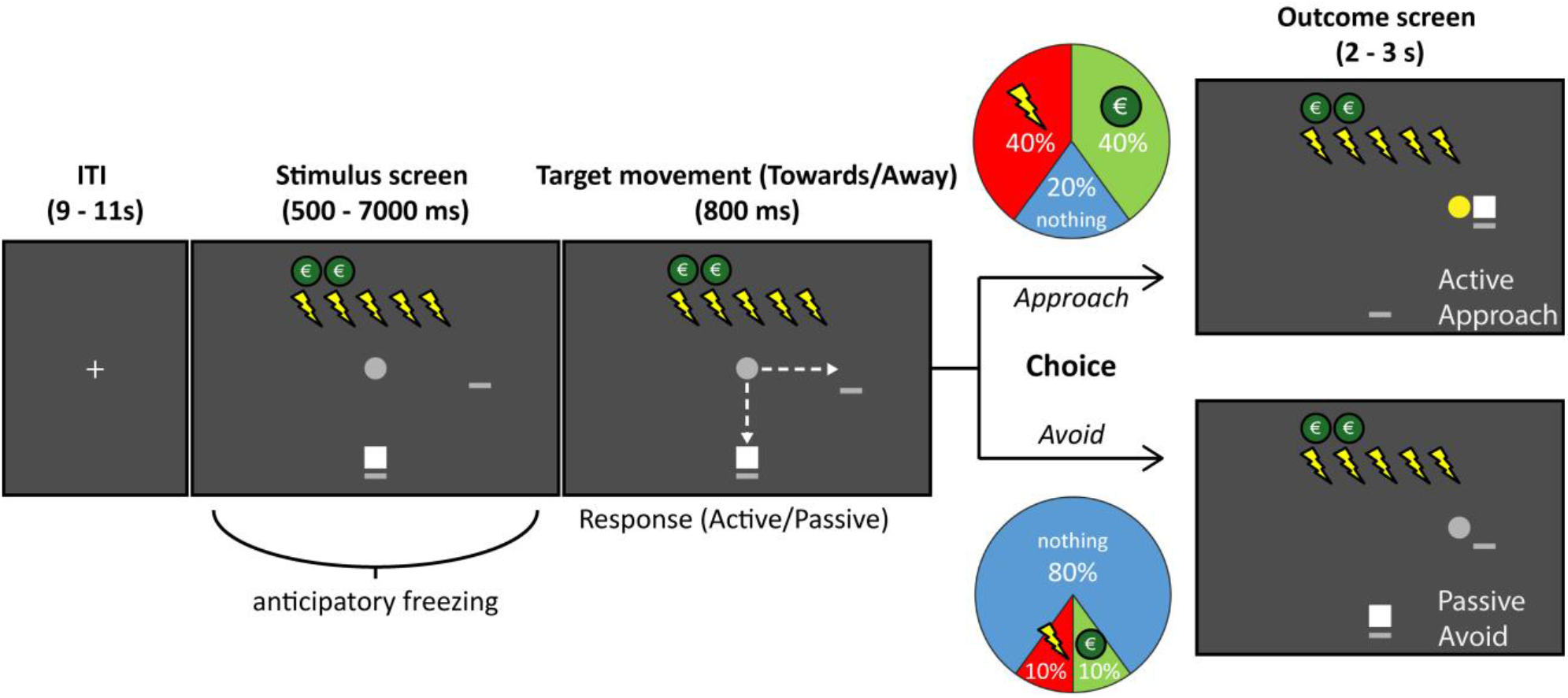
Timeline of an example trial in the Passive-active Approach-avoidance Task (PAT). After presentation of the trial-specific potential money and shock amounts (in this example 5 shocks and 2 Euro), the participant (white square) is asked to approach or avoid the target (gray circle) as it gradually moves either toward or away from the player (respectively passive vs. active action context, shown here = active). At the end of the trial, the participant probabilistically received either shocks, money, or no outcome. Approach was associated with a high chance of receiving the shocks (40%) or the money (40%), and small chance of receiving nothing (20%), while avoid responses led to a high chance of receiving nothing (80%) and a low chance of receiving either the money (10%) or shocks (10%; see colored pie charts). After the target movement, the outcome of the trial was also indicated by a change of the target color into either green (money) or yellow (shocks), or no change (no outcome), and shocks were paid out immediately during that color change. Note: The dashed arrow lines were not present in the actual experiment.

## 2. Results

### 2.1 Task effects on passive and active approach-avoidance decision-making

Overall, the PAT showed the expected effects of potential money and shock amounts on choice behavior. A Bayesian generalized mixed-effects model showed that participants approached the targets significantly more as the amount of money that could be earned increased, and avoided significantly more as the number of shocks that could be received increased (B_money_ = 1.41, 95% CI = [1.17, 1.70], *pp_<0_* < .001; B_shocks_ = −1.08, 95% CI = [−1.38, −0.80], *pp_>0_* < .001; note that effects were considered significant if the 95% CI did not include 0, which corresponds to a posterior probability (*pp*) value < .025, see **Methods**). Moreover, the effects of the amount of money and shocks on choice behavior interacted significantly (B_money:shocks_ = 0.33, 95% CI = [0.15, 0.51], *pp_<0_* < .001). Thus, with increasing amounts of potential money the effect of the shocks on avoidance behavior reduced, and with lower amounts of money the avoidance-inducing effects of high threat increased. Together, these results show that our new task is successful in evoking trade-offs between varying amounts of money and shocks (**Figure 2A**).

**Figure 2.**
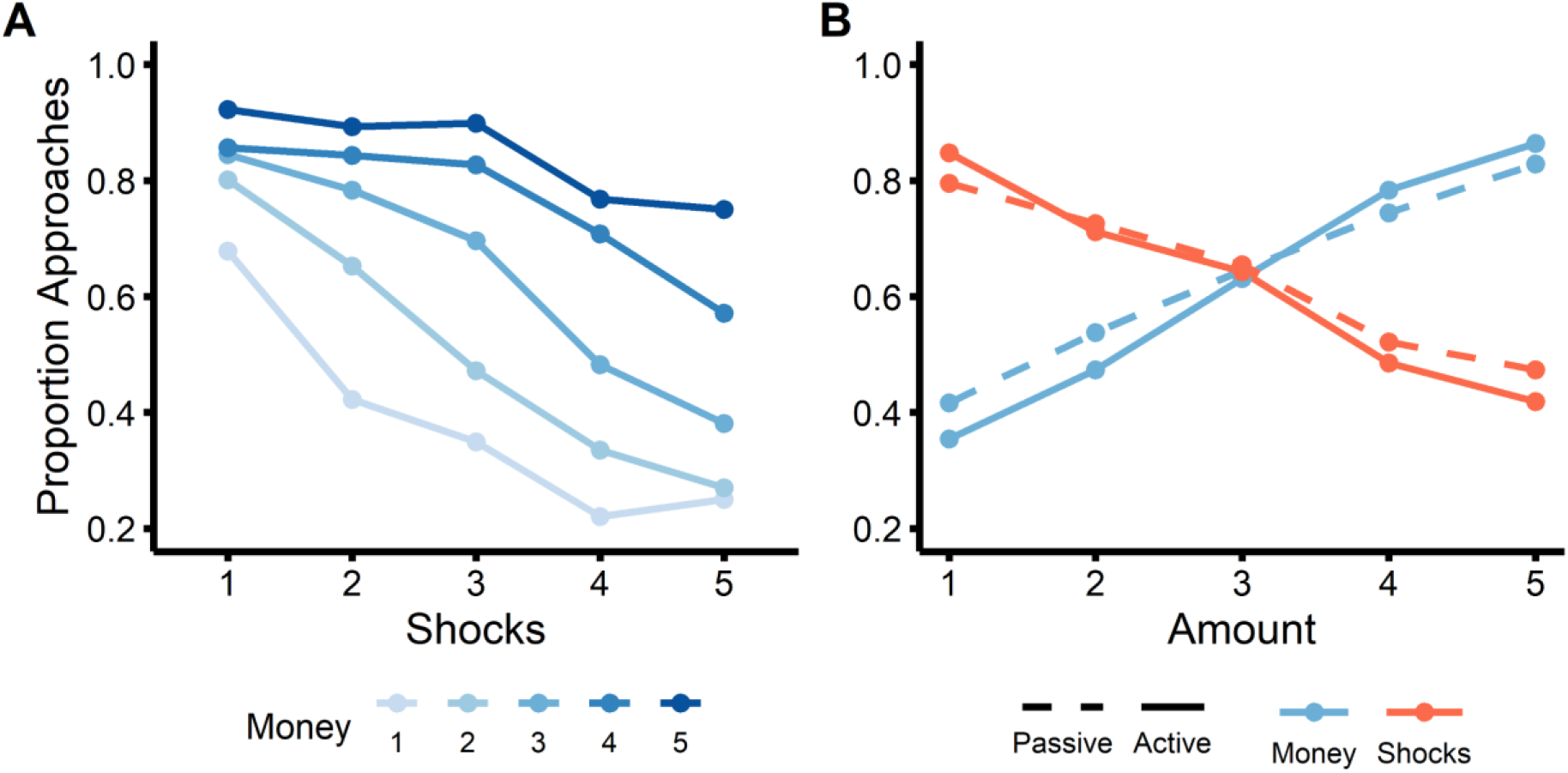
Effects of potential money and shocks and passive/active trial context on the proportion of approach choices. **A)** Participants approach more with higher money amounts, and approach less with higher shock amounts. **B)** Moreover, approach choices are more often active with increased money amounts and more often passive with higher shock amounts. Line types reflect active (solid) vs. passive (dashed) approach conditions.

Importantly, while there were no main effects of action context on approach-avoidance choices (**Figure 2B**; B_actioncontext_ = −0.11, 95% CI = [-0.33, 0.10]), an interaction effect with the potential money and shock amounts showed that the effect of the amount of money and shocks on choice behavior was smaller in the passive action context compared to the active action context (B_actioncontext:money_ = −0.33, 95% CI = [−0.47, − 0.20]; B_actioncontext:shocks_ = 0.26, 95% CI = [0.15, 0.38]). More specifically, higher money amounts increased approach choices more strongly in active approach trials than passive approach trials, while higher shock amounts increased avoidance choices more strongly in passive compared to active trials (**Figure 2B**). This appeared to be driven by a significant invigoration of active responses by increased money amounts, and by an inhibition of action by higher shock amounts (see **Supplementary Results** for details). A similar pattern of action invigoration and inhibition by money and shocks, respectively, was observed in the response times (see **Supplementary Figure 1**). Overall, while we had no explicit expectations on the effect of reward and punishment on action, these findings suggest that increasing potential reward invigorates active compared to passive choices whereas increasing threat leads to more passive (compared to active) choices.

### 2.2 Psychophysiological dynamics of decision-anticipatory freezing

Next, we verified whether our threat manipulation (i.e., number of shocks) impacted freezing-related bradycardia in the decision-anticipatory time window. Therefore, we ran a separate Bayesian generalized mixed-effects model with trial-by-trial heart rate (HR) as dependent variable and continuous predictors money, shocks, their interaction, and body sway (BS). The latter measure was added to confirm the relationship between bradycardia with body sway as an index of freezing in the context of our novel value-based decision task. Here, we indeed observed a significant general deceleration of the average heart rate signal during the anticipation phase relative to the 1s pre-trial baseline (**Figure 3A**, B_intercept_ = −1.01, 95% CI = [−1.73, −0.31], *pp*_>0_ < .003). This deceleration was not further moderated by money (95% CI = [-0.09, 0.43], *pp*_<0_ = .1) or shock amounts (95% CI = [-0.38, 0.19], *pp*_>0_ = .25). Importantly, heart rate was positively related to body sway, confirming a trial-by-trial correlation between anticipatory bradycardia and movement cessation also found in previous freezing literature (Azevedo et al., 2005; Hashemi, Zhang, et al., 2019; Niermann, Figner, Tyborowska, Cillessen, & Roelofs, 2018; Roelofs, Hagenaars, & Stins, 2010) (B_BS_ = 0.61, 95% CI = [0.81, 1.08], *pp*_<0_ < .003). To further explore the effect of the varying shock amounts on heart rate signal dynamics, in a data-driven approach we tested whether the steepness of the slope of the heart rate between the initial peak (±3s) and subsequent trough (±8s) was affected by the number of shocks. Indeed, a mixed model on the slope revealed that higher potential shock amounts led to steeper negative slopes in the heart rate signal (i.e., stronger deceleration; B_HRslope_ = −0.37, 95% CI = [−0.53, −0.02], *pp*_>0_ = .018). This, together with increased skin conductance responses as a function of increased shock amounts (see **Supplementary Figure 5**), indicated that participants were psychophysiologically affected by our threat manipulation.

**Figure 3.**
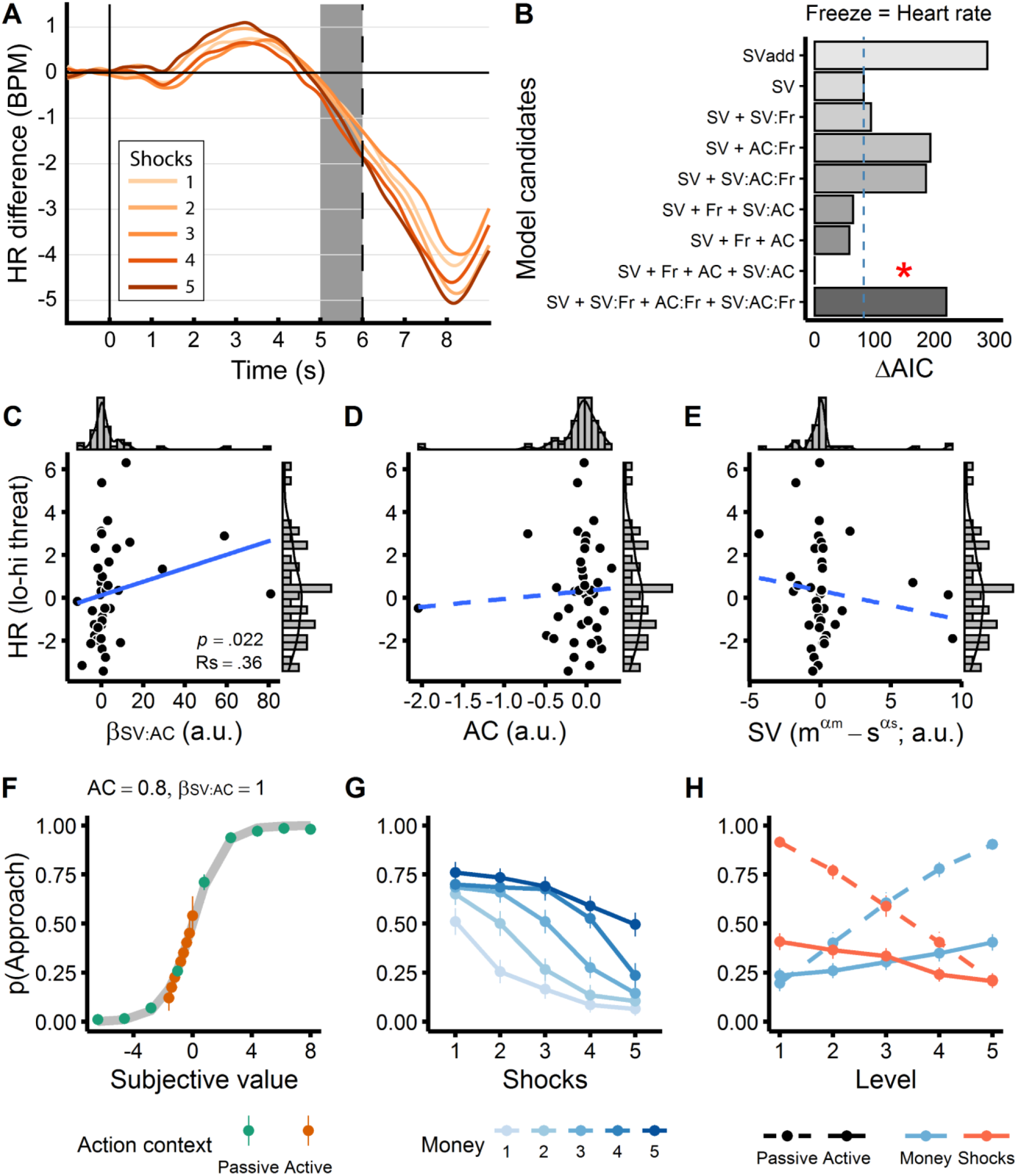
Heart rate and computational modeling results. Panel **A** depicts the average trial time course of baseline corrected heart rate (HR) as a function of the varying shock amounts, with the gray shaded area representing the time window of interest used in all analyses. The vertical dashed line represents the earliest possible target movement onset. Model comparison of various freeze models (**B**) showed that the best fitting model included the subjective value (i.e., potential money and shock amounts), freezing, the action context, and the interaction between the subjective value and action context parameters (indicated with a red asterisk). Additionally, on the subject level, stronger threat-related bradycardia was related to the integration of action context (AC) and subjective value (SV) for approach-avoidance choices (**C**), but not with AC or SV alone (**D, E**). Simulations show that higher values of this interaction parameter led to a diminished effect of subjective value on choice in situations incongruent with passive/active response tendencies (**F, G, H;** for brevity, only simulations of passive ‘participants’ with a positive SV:AC interaction are shown here, see **Supplemental Information** for the full results). Model AIC scores are plotted as the difference from the best fitting model (ΔAIC); lower AIC scores indicate better fit. The blue dashed line (**B**) represents the reference model’s fit and the red asterisk indicates the overall winning model. Solid vs. dashed regression lines in **C-E** reflect significant vs. non-significant relationships. Error bars in **F-H** represent one standard error of the mean (SEM). SV = subjective value, AC = action context, Fr = umbrella term for freeze measures: All models were fitted with either heart rate or body sway as freeze index; see **Supplementary Information**; colons (:) denote interactions.

Interestingly, and in line with previous studies showing action preparation during freezing (Hashemi, Gladwin, et al., 2019), trial-by-trial analyses indicated that stronger bradycardia was associated with faster responding regardless of choice (**Supplementary Figure 2**; B_HR_ = 0.03, 95% CI = [0.01, 0.06], *pp*_<0_ < .006). In addition, we found no relationship of heart rate with approach-avoidance choices or passive vs. active responding, suggesting that - contrary to our hypotheses - trial-by-trial freeze-related bradycardia is not directly related to increased avoidance or passive vs. active responding.

### 2.3 Computational modelling of freezing in approach-avoidance decision-making

Next, we used computational modeling to investigate whether freezing interacts with the subjective value of the potential outcome of approach-avoidance choices (SV) or with the action context (AC). Subjective value reflects the subjective influence of potential money and shock amounts on approach-avoidance choices, whereas the action context parameter AC makes use of the action contexts to capture an individuals’ general tendency to respond passively (AC>0) or actively (AC<0). We performed model comparison on various model candidates (**Table 1**) to investigate how freezing (i.e., heart rate) is most likely to interact with approach-avoidance choices.

**Table 1.** Name and parametrization of all candidate models

|  | Model name | Extra parameters relative to the reference model | Number of free parameters |
| --- | --- | --- | --- |
| 1 | SV<sub>add</sub> | (no $\beta_{m:s}$) | 2 |
| 2 | SV<sup>†</sup> | - | 3 |
| 3 | SV + SV:Fr | $\beta_{SV:Fr}$ | 4 |
| 4 | SV + AC:Fr | $\beta_{AC:Fr}$, AC | 5 |
| 5 | SV + SV:AC:Fr | $\beta_{SV:AC:Fr}$, AC | 5 |
| 6 | SV + Fr + SV:AC | $\beta_{Fr}$, $\beta_{SV:AC}$, AC | 6 |
| 7 | SV + Fr + AC | $\beta_{Fr}$, AC | 5 |
| 8 | SV + Fr + AC + SV:AC | $\beta_{Fr}$, AC, $\beta_{SV:AC}$ | 6 |
| 9 | SV + SV:Fr + AC:Fr + SV:AC:Fr | $\beta_{SV:Fr}$, $\beta_{AC:Fr}$, $\beta_{SV:AC:Fr}$, AC | 7 |

*Note.* m = money; s = shocks; AC = action context; SV = subjective value, Fr = umbrella term for freeze measures: each model was fitted once with heart rate (HR) and once with body sway (BS; reported in supplements); SV<sub>add</sub> = additive SV model; <sup>†</sup>reference model; colons (:) denote interactions

This revealed that the model including the subjective value of the potential outcomes, freeze-related heart rate (as a main effect), the action context parameter AC, and the interaction between the subjective value and the action context parameter (SV:AC) yielded the best fitting model (i.e., lowest AIC; red asterisk in **Figure 3B**). Indeed, although we found no average main effect of passive-active trial contexts on choice behavior, accounting for individual differences in passive/active action tendencies and their interaction with the subjective value of money and shock amounts allows us to better explain approach-avoidance choices. As a check we performed the same model comparison with body sway as alternative freezing index which yielded the similar results and conclusions (see **Supplementary Figure 3**).

While these model comparison findings provide some insight into how general heart rate indices of freezing are most likely to play a role in the decision-making process on the trial-by-trial level, the question remained whether threat-related freezing on the subject level would interact with the subjective value of the choice or the action context. Is the extent to which a person freezes in response to threat related to how they resolve approach-avoidance conflicts? According to our hypotheses, freezing could interact with subjective value (m^αm^ - s^αs^) or action implementation alone (AC parameter), but we also considered the possibility that it relates to how the subjective value *interacts* with passive versus active responses (β_sv:Ac_). To test this, we performed exploratory correlations on the threat-induced individual differences in bradycardia and the parameter estimates of the winning computational model (**Figure 3B**). This threat-related bradycardia did not correlate to the action parameter AC or subjective value alone (both *p* > .1), but did correlate with their interaction (β_sv:Ac,_ Rs = .36, *p* = .022, uncorrected for 3 correlations; **Figure 3C-E**).

To gain further understanding of the positive relationship between threat-related bradycardia and the β_sv:Ac_ parameter, we performed simulations of approach-avoidance choices for positive vs. negative values of the β_sv:Ac_ parameter, both for ‘participants’ with passive and active response tendencies. These revealed that more positive βsv:Ac parameter values result in a diminished effect of potential money and shock amounts on choice behavior for active approach trials in passive participants (AC>0) and for passive approach trials in active participants (AC<0), and vice versa for negative β_sv:Ac_ values (**Figure 3F-H** and **Supplementary Figure 4**). More generally, these simulations indicate that individuals with stronger freeze-related bradycardia may differentially weigh the influence of potential reward and threat outcomes depending on the action context. When the action context does not match the internal action tendency, stronger freezing is associated with a diminished effect of subjective value on choice, whereas in congruent action contexts the effect of subjective value on choice remains relatively unaffected.

## 3. Discussion

This study aimed to bridge the fields of affective and decision sciences by systematically manipulating psychophysiological threat-states and action contexts during approach-avoidance conflict resolution. Three main findings contribute novel insights into decision making under acute threat. First, at the behavioral level, participants were well able to trade-off money versus shocks and whereas high threat was associated with more passive responding, high reward induced more active responding. Interestingly, this motivation-related invigoration of action was also observed in the response times. Second, freezing-related bradycardia was related to faster responding but there was no direct relationship with choice or action. Third, approach-avoidance decisions were best predicted by a model that included not only the subjective value of the potential outcomes, but also incorporated bradycardia and the action context. Moreover, at the subject level, threat-related bradycardia was positively correlated to the interaction between the action (passive/active) parameter and the subjective value. Specifically, high freezers show a diminished effect of subjective value on choice when the action context does not match their individual passive/active action tendencies, signifying a potential relationship between freezing and the way decisions are made on the subject level. Together, these findings show that integration of psychophysiological states as well as action context into our decision models, yields new mechanistic insight into how approach-avoidance conflict decisions are made under threat. This has implications for behavior in real-life threatening situations, such as for first-responders or anxiety patients, suggesting that it may be beneficial to learn to modulate freezing-related states (e.g., using bio-feedback (P. Lehrer et al., 2020; P. M. Lehrer & Gevirtz, 2014)) to help optimally weighing values in the decision. Given this relevance of approach-avoidance conflict decision-making for real-life threatening situations, these findings call for models of value-based decision-making that better integrate internal (e.g., value, psychophysiology, & action tendencies) and external (e.g., action context) components of the decision.

Anticipation of decisions was associated with freezing-like reactions, both in terms of heart rate deceleration (bradycardia) and body sway reduction. Following previous work (Azevedo et al., 2005; Hashemi, Zhang, et al., 2019; Niermann et al., 2018; Roelofs et al., 2010) and relevant for subsequent neuroimaging studies, we validated the heart rate measure showing a correlation to a concurrent bodily freezing measure. Freezing-related bradycardia did not interact with approach-avoidance conflict decisions directly, but seemed to rather act on the implementation of the required action as well as its interaction with subjective value.

Regarding the relation between freezing and action implementation, freezing-related bradycardia was associated with significantly faster response times for subsequent action, supporting a role of freezing in action preparation. Although few previous animal studies suggested freezing to be part of a general response inhibition (Bolles, 1970; Martinez et al., 2013; Moscarello & LeDoux, 2013), our results are more in line with recent human literature consequently showing that across parameters (i.e., heart rate, body-sway, and neural (midbrain PAG) activity), increased freezing magnitude is associated with faster subsequent action execution, without going at the expense of an increase in error rates (Gladwin et al., 2016; Hashemi, Gladwin, et al., 2019). This discrepancy is likely related to the fact that freezing in the animal literature is typically defined as the reduction or absence of movement (Moscarello & LeDoux, 2013). The lack of bradycardia indices makes it harder to distinguish it from other passive defensive states: such as orienting and tonic and collapsed immobility (Kozlowska, Walker, McLean, & Carrive, 2015; Mobbs, Headley, Ding, & Dayan, 2020). Recent animal work that considered heart rate indeed found that bradycardia was related to immobility in anticipation of threat (to a conditioned cue) and not after (unconditioned) threat exposure(Schipper et al., 2019). Also, in the animal literature freezing is often assessed in terms of response duration rather than magnitude of state-changes, confounding the relation with subsequent action. Future research using more precise translation between animal and human paradigms has the promise to assess similarities and discrepancies in the function of freezing across species (Fendt et al., 2020).

Importantly, individual approach-avoidance choices were best predicted by a model including not only the effects of the subjective value of the potential outcomes, but also freeze-related bradycardia and the action context (active-vs-passive). This is in line with the notion that freezing plays a role in decision-making under threat (Blanchard et al., 2011; Ly et al., 2014; Roelofs, 2017), though on the trial level this does not seem to take place through interaction with outcome value computations or action tendencies. Interestingly, on the subject level, the relation between freezing and subjective value depended on the congruency between the passive-active trial context and the subject’s passive-active response tendency. Specifically, stronger threat-related bradycardia was associated with a diminished effect of subjective value on choice in situations that are incongruent with their passive or active response tendency. This finding may be interpreted in line with previous notions that people with stronger threat-induced freezing show general upregulation of sensory systems (Lojowska et al., 2018) and better risk assessment (Blanchard et al., 2011). Such upregulation may be hampered when counterintuitive actions have to be prepared. Speculatively, contexts that do not match some internal action tendency may put increased demand on cognitive control processes involved in allocation of resources needed to weigh decision-relevant information (Bramson et al., 2020; Eder & Hommel, 2013; Frijda, Ridderinkhof, & Rietveld, 2014; Ridderinkhof, 2017).

The notion that individual bias in passive versus active responding matters in the relation between freezing and subjective value may explain that the results from our mixed effect models showed no *average* trial-by-trial relation between freezing and approach-avoidance choices. Likewise, Ly and colleagues (Ly et al., 2014) found a relation between freezing and subsequent approach-avoidance decisions between-subjects but not within. Together, the between-subject nature of our freezing and action-context findings may also imply that our freezing effects may also partly reflect a trait-like effect, an interpretation that is further supported by genetic components previously identified for individual differences in freezing-tendencies (Niermann et al., 2019; Schipper et al., 2019).

The relevance of considering active versus passive decisions during approach-avoidance conflict was not only supported by its interaction with bradycardia. Our computational modeling approach showed that the freeze model that accounted for individuals’ passive-active response tendencies as well as its interaction with the subjective value of the outcomes, explained choice behavior significantly better than a standard subjective value model. Finally, the relevance of taking action context into account was supported in our task-validation mixed effects models approach, where it helped to reveal a clear pattern of reward-induced action invigoration: higher money amounts led to more active responses, whereas more potential shocks led to more passive responses. These findings fall in line with many studies in the go/no-go literature that demonstrate action invigoration and inhibition in the face of reward vs. threat (Guitart-Masip, Duzel, Dolan, & Dayan, 2014; Guitart-Masip et al., 2012). Mimicking these choice effects, we additionally observed faster response times for approach choices for increased money amounts, and faster responses for avoid choices for increased shock amounts, suggesting motivation-related invigoration of action. Together, these finding show that it is relevant to incorporate psychophysiological states and action context into value-based models of approach-avoidance conflict decisions.

Some limitations should be considered when interpreting these findings. First, although freezing temporally preceded the choice behavior in our paradigm, we cannot make *causal* inferences on the role of freezing in decision-making. Such causality may be achieved in future studies, for example by stimulating the vagal nerve or by stimulating brain regions that are thought to be causally involved in the regulation of freezing-related bradycardia, such as deep brain stimulation of the PAG (A. L. Green et al., 2006; Sims-Williams et al., 2017). Second, although this is the first study testing approach-avoidance conflict resolution under relatively ecologically valid conditions of acute and primarily reinforced threat (threat-states being validated in all three psychophysiological measures), follow-up studies using ambulatory monitoring of psychophysiological states should test whether findings generalize to real-life decisions. Third, because the interpretations of the correlation between threat-related bradycardia and model parameters are based on an exploratory analysis, they should be regarded as hypothesis-generating rather than hypothesis-testing. Still, while provisional, these findings provide deeper insight into the relationship between freezing and decision-making which could be meaningfully interpreted using model simulations.

In conclusion, we present a novel paradigm in which we could capture how threat-related psychophysiological states such as freezing-related bradycardia are associated with passive and active approach-avoidance conflict-resolution. Our results support the relevance of considering individuals’ tendency to respond passively versus actively and the role of such action-related parameters in explaining choice behavior besides the subjective value associated with approach and avoidance. Second, although bradycardia may not be related to changes in approach-avoidance conflict decisions by directly affecting the subjective value of the potential outcomes, it interacts with *how* approach-avoidance conflict decisions are made on the individual level, coinciding with a diminished effect of the subjective value of the choice outcomes in situations that are incongruent with the dominant passive or active response tendency. Critically, our findings illustrate that it is important to incorporate concurrent *external* action contexts as well as *internal* psychophysiological states of the decider in models of approach-avoidance conflicts.

## 4. Methods

This study was preregistered on the Open Science Framework before data collection (https://osf.io/68e75). All research activities were carried out in accordance with the Declaration of Helsinki and approved by the local ethics committee (Ethical Reviewing Board CMO/METC [Institutional Research Review Board] Arnhem-Nijmegen, CMO 2014/288) and conducted according to these guidelines and regulations (i.e., medical/scientific research).

### 4.1 Participants

A total of 42 participants (aged between 18 and 35 [*M±SD* = 23.5±3.63]; 24 females) were included in the study. An additional three participants were excluded due to technical issues (n=2) and dizziness (n=1). Sample size was determined using a simulation-based power analysis (pilot sample of n=5) which revealed a sample size of 40 participants was sufficient to detect a small (i.e., odds ratio of 1.5(Chen, Cohen, & Chen, 2010)) heart rate by shock interaction effect with at least 80% power (*simr* package in R (P. Green & McLeod, 2016; R Core Team, 2019; RStudio Team, 2019)). Inclusion criteria were age (younger than 18 or older than 35), Dutch or English speaking, and right-handedness; exclusion criteria were self-reported current pregnancy, current or lifetime history of psychiatric, neurological, or cardiovascular disorder, endocrine illness, claustrophobia, plaster allergy, and self-reported high or low blood pressure. All participants gave written informed consent before participation, and were paid for participation (€16) plus bonus money contingent on their task choices (see below).

### 4.2 Experimental procedure

Participants came to the lab once for a 90 min. assessment involving the Passive-active Approach-avoidance Task (PAT). Procedures started with performing a standardized 5-step shock work-up to calibrate the intensity of the electric shock which remained the same for the rest of the task (see **Peripheral stimulation and measurements**).

Next, the participant stepped on the stabilometric force platform, read through the onscreen instructions, and performed practice trials. After the experimenter verified that instructions were understood, participants started the PAT paradigm which consisted of 150 trials divided up in 10 blocks (±5 min duration). Participants were encouraged to take breaks by stepping off the platform in between blocks to minimize the chance of exhaustion and potential dizziness due to prolonged standing still.

Afterwards, participants received one more electric shock and were asked to rate the intensity. Finally, we administered the State Trait Anxiety Questionnaire (STAI (Spielberger, Gorsuch, Lushene, Vagg, & Jacobs, 1983); *M±SD* = 33.94±8.84) and the Beck Depression Inventory II (BDI-II (Beck, Steer, & Brown, 1996); *M±SD* = 6.38±4.92) to characterize the sample.

### 4.3 Experimental Paradigm - Passive-active Approach-avoidance Task (PAT)

In the Passive-active Approach-avoidance Task (see **Figure 1**), participants were instructed to make approach-avoidance decisions in response to a moving target that was associated with varying amounts of monetary rewards and shocks. On each trial, they either approached or avoided the target by positioning the player icon towards or away from the target, which probabilistically led to either money, electric shocks, or no outcome. Since extensive pilot work had indicated that individuals were well able to trade-off money (1-5 Euro) and shocks (1-5), we did not compute subject-specific money-to-shock indifference pairs; all participants received the same monetary offers. The monetary outcome of three randomly selected trials (max. €15) was paid out as a bonus fee. Finally, to be able to distinguish between active and passive approach-avoidance decisions, responses were given in two possible action context conditions (passive/active). This was determined by the movement direction of the target, which was either towards or away from the player icon. If the target moved away from the participant an approach decision involved an active response and avoid was passive. In case the target moved towards the participant, avoidance required an active response and approach was passive. Per trial, we recorded the participants’ choice (approach/avoid), response type (passive/active), and response time. Psychophysiological measures of heart rate, body sway (i.e., postural freezing), and skin conductance (sympathetic control) were recorded throughout the task. For a detailed description of the trial procedure, see **Supplementary Information**.

### 4.4 Peripheral stimulation and measurement

Electric shocks were delivered using a 9V MAXTENS 2000 shocker machine (Bio-Protech inc., 2019) and standard Ag/AgCl electrodes which were attached to the distal phalanges of the fourth and fifth fingers of the left hand. Electrocardiogram (ECG), bodily displacement, and electrodermal activity were recorded using a BrainAMP EXG MR 16 channel amplifier, an EXG aux device, and BrainVision Recorder software (Brain Products Gmbh, 2019). To measure heart rate, we made an ECG using two standard Ag/AgCl measurement electrodes placed just below the right collar bone and around the lowest left rib (i.e., diagonally through the heart), and a third ground electrode positioned just below the left collar bone (i.e., above the heart). We measured body sway by having participants stand on an in-house built 1×1m stabilometric force platform that detects the participant’s displacement relative to the center of pressure, or body sway, in the anterior-posterior and medio-lateral direction. Electrodermal activity was assessed using two Ag/AgCl electrodes attached to the second and third distal phalanges of the left hand with some non-adhesive gel to improve signal quality. Skin conductance responses were only used to account for sympathetic activity in control analyses. Finally, because of the slow development of the heart rate signal over time, only trials with a long stimulus screen duration (i.e., ≥6 sec, 100 trials per participant) were included in data analysis. See **Supplementary Information** for a more detailed description.

### 4.5 Data analysis

#### 4.5.1 Quantification of freeze measures

For our analyses we computed trial-by-trial quantifications of our psychophysiological indices of freezing (i.e., heart rate and verified by body sway) by extracting per trial the mean (HR) or median (BS) signal over a time window of 5000-6000ms after trial onset (i.e., right before the earliest possible target movement onset). See **Supplementary Information** for details.

#### 4.5.2 Behavioral analysis

For our behavioral analyses, we complemented our planned computational modelling approaches with mixed-effects modeling to validate our novel approach-avoidance conflict task. Mixed-effects models are well suited to robustly estimate and statistically test average (fixed) effects, while still controlling for individual differences by estimating random effects. Computational modeling (see section below) was used to perform model comparison of different models that included freezing as a parameter. Additionally, parameter estimates of the winning model were correlated to threat-induced bradycardia. All mixed and computational modeling results reported in the main text were robust after controlling for sympathetic activity (i.e., skin conductance), depression and anxiety scores (i.e., BDI-II (Beck et al., 1996) and STAI (Spielberger et al., 1983)), and gender (binary; see **Supplementary Information**).

We ran three separate (generalized) Bayesian mixed-effects models to test task effects on choice [approach/avoid], response type [passive/active], and response times (see **Table 2** for all predictors per model; *brms* package in R (Bates, Mächler, Bolker, & Walker, 2015; Bürkner, 2017, 2018; R Core Team, 2019; RStudio Team, 2019)). Because Bayesian analyses do not yield traditional *p*-values, effects were considered ‘significant’ if the 95% credible interval (CI) of the posterior distributions did not include 0. We additionally report *pp* (i.e., posterior probability) values which are defined as the proportion of the posterior parameter distributions that lies above or below 0 (depending on the direction of the effect) (Maier, Raja Beharelle, Polanía, Ruff, & Hare, 2020; Polania, Woodford, & Ruff, 2018). To be able to interpret them as two-tailed tests that directly reflect the significance of the 95% CI, *pp*-values are significant when they are smaller than 0.025 (Makowski, Ben-Shachar, Chen, & Lüdecke, 2019). For all parameter estimates we used the default *brms* priors. All Bayesian mixed-effects models followed a maximal random effects structure unless otherwise specified (i.e., by-participant random intercepts, by-participant random slopes for all within subject effects, and random correlations for all combinations of random intercepts and slopes (Barr, Levy, Scheepers, & Tily, 2013); see **Supplementary Information** for details on priors and model estimation settings).

**Table 2.** Model specifications of mixed models to test task-effects on choice behavior and psychophysiology

| Dependent variable | Categorical predictors | Continuous predictors |
| --- | --- | --- |
| Choice (approach/avoid) | Action context (passive/active) | Money; Shocks; Heart rate |
| Response type (passive/active) | Choice (approach/avoid) | Money; Shocks; Heart rate |
| Response time | Choice (approach/avoid) | Money; Shocks; Heart rate |
| Heart rate | - | Money; Shocks; Body Sway |

*Note.* Money and shock amounts ranged 1 – 5.

#### 4.5.3 Computational model

The effect of trial-by-trial potential outcomes (i.e., money and shock amounts) on approach-avoidance choices was formalized according to the following decision value function:

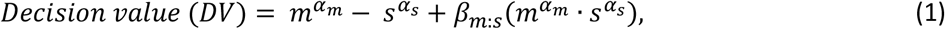

with exponential free parameters α_m_ and α_s_ modulating the effect of the varying money (m) and shock (s) amounts, and free parameter β_m:s_ weighting their interaction. Since the decision value function in equation (1) estimates only the influence of the subjective value (SV) of the potential outcomes on choices without taking into account freezing and the action context, it is further referred to as the SV model (Park et al., 2011). To test whether freezing interacted with the subjective value of the choice or with the action context, this model was then extended via stepwise inclusion of additional parameters of interest that capture the contribution of psychophysiological freezing responses (heart rate measure weighted by multiplicative parameter β_HR_), individuals’ passive-active action tendencies (action context parameter AC), and their interactions (see **Table 1**). For simplicity, in interaction terms the subjective value of outcomes was operationalized as simply the difference between the money and shock levels (*m^a_m_^ — s^a_s_^*). The interaction between, for example, the subjective value and action context parameter AC is thus formalized as (*m^α_m_^ — s^α_s_^*)·AC.

The action parameter AC is defined as

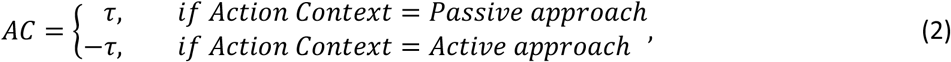

and it captures the variance in choice behavior related to active vs. passive responding. A positive AC parameter indicates a stronger tendency for passive responses, whereas a negative AC indicates a tendency to respond actively.

To address our hypotheses, our freezing parameter (β_HR_) was included as a main effect and in interaction with the previously mentioned SV and AC terms. Though heart rate was our main index of freezing and body sway was only intended as a validation method, we also fitted the same models with body sway as an alternative freeze index to check robustness of the results (see **Supplementary Results**). Together these two are referred to as ‘freezing’ but are in fact separate parameters estimated in separate models.

The complete decision value function of each model was transformed into a choice probability by passing it through a soft-max logistic choice rule:

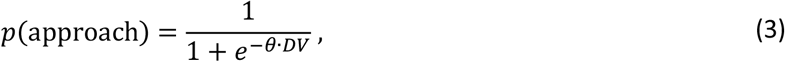

in which θ is a sensitivity parameter that modulates the slope of the decision value function in terms of choice stochasticity. While all models were initially fitted with θ as a free parameter, we re-fitted the models with a representative fixed θ-value of 5 (determined by taking the best fitting model’s average θ across participants) in order to increase model stability and allow more reliable interpretation of the parameter estimates (Gershman, 2016; Stewart, Scheibehenne, & Pachur, 2018). The model comparison results were unaffected by this change.

Based on our hypotheses that freezing interacts with subjective value computations and passive-active action tendencies, we selected candidate models for model comparison that iteratively include parameters that denote main effects of the subjective value and action context, the subjective value-by-action interaction (SV:AC), and freezing (heart rate or body sway) as a main or interaction effect (**Table 1**). Specifically, we first created models that separately included freezing in interaction with the subjective value, action, or the SV:AC interaction (models 3-5 of **Table 1**). Then, we included freezing as a main effect in addition to the subjective value, action, or the SV:AC interaction (models 6-8). Finally, we included a single model in which freezing interacts with all of the subjective value, action, and SV:AC parameters simultaneously (model 9). The SV model as described in equation (1) was our main reference model since it closely resembles relevant pre-existing computational models of approach-avoidance conflict decisions (Park et al., 2011) and still retains parsimoniousness (model 2). All model candidates were fitted and subsequently compared using maximum likelihood estimation (Bolker & R Core Team, 2017; Byrd, Lu, Nocedal, & Zhu, 1995) and the Akaike Information Criterion (Akaike, 1974) (see **Supplementary Information** for details).

Additionally, parameter estimates of the winning model were correlated to a threat-dependent freeze measure (i.e., threat-induced bradycardia) using Spearman rank correlations (alpha of .05) to account for any outlier and non-normality related issues. This was computed as the average difference in heart rate deceleration between low (≤ 2 shocks) and high threat (≥ 4 shocks) conditions, which approximates previous quantifications of freezing that contrast low threat and high threat conditions(Gladwin et al., 2016; Hashemi, Gladwin, et al., 2019). For the correlation of this threat-related bradycardia with the subjective value of the potential outcomes, we computed the average subjective value per participant using median money and shock levels of 3 (different values did not affect the correlation). Finally, simulations were performed to verify plausibility of the predictions of the winning model (see **Supplementary Information**). All computational modeling, correlation, and simulation analyses were performed in R 3.6.1(R Core Team, 2019).

## 5. Acknowledgements

This work was supported by a consolidator grant from the European Research Council (ERC_CoG - 2017_772337) awarded to KR.

## 6. Author Contributions

F.H.K., L.H., B.F., J.X.O., and K.R. conceived and designed the study, F.H.K. and L.H. collected the data, F.H.K. performed the data analysis, all authors were involved in data interpretation, F.H.K. wrote the first full draft, and all authors contributed to the final manuscript.

