## Supplementary Information for "Defensive freezing and its relation to approach-avoidance decision-making under threat"

**Supplementary Methods**

***PAT trial procedure***

Each trial started with a variable inter-trial interval (ITI) of 9-11s period to allow a return to baseline for all psychophysiological measures (Klumpers, Kroes, Baas, & Fernández, 2017). Next, participants were faced with the stimulus screen including the player and target icons, and the corresponding levels of potential money (green €-icons indicating 1 Euro each), and electric shock amounts (lightning bolts indicating 1 shock each). The player icon was a white square and was always positioned in the lower center of the screen. The target was represented by a grey circle positioned in the center of the screen. The icons indicating the money and shock amounts were always presented at the top of the screen and ranged from 1 to 5. For the main analysis there were 25 possible combinations of money and shock levels (5 money x 5 shocks) which were repeated four times (i.e., 100 trials in total). For these trials, the entire stimulus screen was presented for a variable stimulus-to-movement interval (SMI) of 6-7s. This length was chosen because of the slow development of the psychophysiological signals (Gladwin, Hashemi, van Ast, & Roelofs, 2016; Hashemi, Gladwin, et al., 2019; Lojowska, Gladwin, Hermans, & Roelofs, 2015; Lojowska, Ling, Roelofs, & Hermans, 2018). To ensure prolonged anticipation of the upcoming target movement we also included 30 trials with shorter SMIs of 0.5-6s which were not included in the analysis. Lastly, we also included 20 trials with 0-money or 0-shock amounts which served as a manipulation check to assess whether participants were paying attention to the task. Logically, participants should not approach when no reward can be obtained and vice versa for shocks. Indeed, participants avoided 0-money trials and approached 0-shock trials (proportion approaches $M \pm SD_{0\text{-}money}$ = .09%±.14; $M \pm SD_{0\text{-}shocks}$ =.90%±.17). After the SMI, to enable passive vs. active action contexts, the target gradually moved to one of two possible locations: either downward towards the player (passive context) or away from the player icon (i.e., left or right [randomized across trials]; active context). These locations were indicated onscreen by grey placeholder bars. Each action context (i.e., passive and active) made up 50% of the trials (i.e., there were 50 active and 50 passive trials). While the target was moving (duration of 800ms), participants had to either approach or avoid the target by positioning themselves at the same location as the target (i.e., approach) or at the other location (i.e., avoid). Depending on the target movement direction, this required an active response by moving the player icon to the other location (i.e., via a button press), or a passive response by leaving the player icon where it was (i.e., no button press). This led to four different response types: active approach and passive avoid when the target moved away from the player icon (active context) and active avoid and passive approach when the target was moving towards the player icon (passive context). Responses were only recorded as active when they occurred during the target movement phase. Note that not pressing a button during that phase would be recorded as a passive response.

As soon as the target stopped moving, the outcome was indicated by color coding of the target as green (i.e., money), yellow (i.e., shocks) or grey (i.e., no outcome). The task was probabilistic meaning that if the participant approached, there was a 40% chance of receiving money, 40% chance of receiving electric shocks, and a 20% chance of receiving nothing. If they avoided, there was still a 10% chance of receiving money, 10% of receiving electric shocks, and an 80% chance of receiving nothing. The 10% payout chance of money and shocks for avoid choices was added to make the task more threatening and thus increase anticipatory freezing, and the 20% chance of no outcome for approach choices was added to match the outcome uncertainty between the two choice options (i.e., 3 possible outcomes per choice, see **Figure 1**). The electric shocks were administered immediately at the end of each trial, to ensure acute threat. Choices were incentivized by paying out the summed monetary outcome of three randomly selected trials after the experiment.

***Electrical stimulation procedure***

Each shock had a duration of 200ms (consisting of 250μs pulses at 150Hz) and an intensity varying in 10 steps between 0-40V/0-80mA across 500Ω. We used a standardized shock adjustment procedure in which participants subjectively rated the intensity of 5 shocks so that the final shock intensity was experienced as uncomfortable, but not painful (final shock intensity was *M*±*SD* = 4.43±1.81).

***Stabilometric platform***

The platform was calibrated before each participant so that the amount of pressure was evenly distributed among all four sensors. Participants were instructed to move as little as possible and to stand in a stable position with their feet positioned approximately 30cm apart.

***Psychophysiological pre-processing details***

All psychophysiological data were preprocessed and analyzed using Matlab2018b and R (R Core Team, 2019; RStudio Team, 2019; The Mathworks, 2018). The raw electrocardiogram signal was downsampled to 250Hz and filtered with a Butterworth bandpass filter (0.5-10Hz) (Hashemi, Gladwin, et al., 2019; Hashemi, Zhang, et al., 2019). Next, R-Peaks were detected using an in-house built peak detection algorithm, and were visually inspected trial-by-trial and manually corrected if necessary. Trials were excluded from analysis if peaks could not be reliably detected within a time window of 1 second pre-trial onset (baseline window) until 6 seconds after the trial onset (earliest possible onset of target movement in long stimulus-to-movement-interval trials). Short and medium SMI trials were never included. Changes in heart rate were calculated in beats per minute (BPM) based on the beat-to-beat inter-beat intervals, and baseline corrected relative to the average heart rate during the 1 second window before trial onset.

The raw signal of the stabilometric platform was also downsampled to 250Hz and filtered with a Butterworth bandpass filter (0.01-10Hz) (Hashemi, Zhang, et al., 2019). The body sway signal was calculated as the standard deviation of the center of pressure in the anterior-posterior (AP) direction within (overlapping) 1000ms time windows surrounding each sample (Gladwin et al., 2016). This was done for each sample from 1000ms pre-trial onset until 6000ms post-trial onset (i.e., covering the baseline window and the entire anticipation period). The signal of each trial was baseline corrected relative to the 1 second pre-trial window. Since there are no clear pre-existing exclusion criteria for body sway signals and we wanted to exclude as few trials as possible, trials were only excluded if more than 50% of all samples within that trial deviated more than 3SD's from the trial mean. Since this is a relatively conservative exclusion criterion and we still wanted to ensure a reliable signal that's robust to the remaining outliers, we computed the median rather than the mean body sway for later summary statistics.

The raw electrodermal activity signal was low-pass filtered (2Hz) and smoothed using a moving-average filter (window of 0.25s/50 samples). Skin conductance responses (SCRs) were for each trial determined as the largest SCR amplitude within a latency window from 0.5 to 6 seconds after trial onset with a max. base-to-peak rise time of 6 seconds. Trials in which no response was detected were given a SCR value of 0, and responses were square root transformed before statistical analysis.

***Quantification of freeze measures***

Based on heart rate patterns in previous studies (Lojowska et al., 2015), we preregistered to average over a time-window of 2 seconds before the target movement initiation for each trial. Inspection of the actual data however indicated we had to make the following minor adaptation. When considering the time window based on each trial specific target movement onset, we realized that in this specific task the trial-by-trial extracted signal could potentially be confounded by the delay of the movement onset (e.g., a lower average heart rate could be due to a delayed movement onset in that trial instead of stronger freezing). This is dealt with by setting it to a fixed post-trial onset time window for all trials (Hashemi, Gladwin, et al., 2019; Klumpers et al., 2017). Previous studies most similar to ours showed most robust threat effects on heart rate and body sway during the very final stage of anticipation, i.e., after the 5 second mark (Hashemi, Gladwin, et al., 2019; Hashemi, Zhang, et al., 2019). Therefore, we decided the 5-6s window provides a more precise index of freeze-related heart rate deceleration and body sway reduction.

***Behavioral analysis – mixed model specifications***

For the Bayesian mixed-effect models, all continuous predictors were zero-centered and scaled, and categorical predictors were coded using sum-to-zero contrasts (Fox & Weisberg, 2019). The freezing indices heart rate (HR) and body sway (BS) were entered into the models as continuous predictors that interacted with all other main-effects and two-way interactions apart from any effect that involved the other freezing index (i.e., HR never interacted with any effect that involved BS, and vice versa). Dependent variables choice (approach/avoid) and response type (active/passive) were modelled with Bernoulli distributions (logit link), and response times were modelled using a shifted lognormal distribution (identity link).

We used the default *brms* priors which are improper flat priors for population-level (i.e., fixed) effects, weakly informative Student-*t* priors for group-level effects (i.e., random intercepts and slopes), and LKJ-Correlation priors for random correlations (Bürkner, 2017, 2018). All models were fit with 6000 samples (3000 warm-up) across 6 chains, of which all converged to a solution without warnings (i.e., all R-hats between 0.99 and 1.01).

***Computational modeling - fitting and comparison procedures***

Computational models were fitted per participant using maximum likelihood estimation and the L-BFGS-B optimization method to allow for box constraints on free parameters (Bolker & R Core Team, 2017; Byrd, Lu, Nocedal, & Zhu, 1995). To prevent converging to local minima in the parameter space, we repeated the fitting procedure for 1500 iterations per participant with starting parameters randomly sampled from normal distributions (i.e., $\mathrm{N}(0, 4^2)$ and $\mathrm{N}(1, 2^2)$ for multiplicative and exponential parameters respectively) and selected the best fitting iteration for each model and participant based on the AIC (Akaike Information Criterion(Akaike, 1974)). Parameter boundaries were set at -100 to 100 for all free parameters (except $\theta$, which had a lower bound of 0.001 before being fixed). The fitted parameter estimates of the best fitting model (for both heart rate and body sway) did not contain any boundary solutions (see **Supplementary Table S1**).

To compare candidate models across participants, we computed per model the sum of the individual model fits across participants. Like in the model fitting procedure, we used the AIC as a goodness-of-fit measure because the BIC (Bayesian Information Criterion (Schwartz, 1978)) assumes that the real data-generating model is in the candidate set (Wagenmakers & Farrell, 2004), which is not an assumption we want to make. Two additional participants were excluded from model comparison because of a strong lack of variability in their choice data (i.e., fewer than 5 avoid or approach choices) which may result in improper parameter estimates, leaving 40 participants.

### Supplementary Results

#### *Supplementary behavioral results*

To test the action invigoration effect observed in main text **Figure 2B** we conducted a second Bayesian mixed model, directly investigating active versus passive responses (dependent variable) as a function of choice, money, shocks, and heart rate. In this model we confirm a significant increase in active responses for greater money amounts ($B_{money}$ = 0.11, 95% CI = [0.04, 0.19], $pp_{<0}$ = .002) and a significant increase in passive responses for increased shock amounts ($B_{shocks}$ = -0.09, 95% CI = [-0.16, -0.02], $pp_{>0}$ = .007). There were no additional interaction effects, nor any effects of heart rate. Testing task effects on response times (RT; active responses only), a Bayesian mixed-effects model with categorical predictor choice (approach/avoid) and continuous predictors money, shocks, and heart rate revealed a marginally significant effect of choice on RT, showing on average marginally faster RTs for avoid compared to approach choices ($B_{choice}$ = -0.03, 95% CI = [-0.066, 0.001], $pp_{>0}$ = .028; remember that $\alpha$ = .025). This relationship was further moderated separately by money and shock amounts: participants became faster for approach decisions as the amount of money increased, but faster for avoid decisions as the number of shocks increased, suggesting motivation-related action invigoration by reward and threat (**Supplementary Figure S1A & B**, $B_{choice:money}$ = 0.04, 95% CI = [0.02, 0.07], $pp_{<0}$ < .002; $B_{choice:shocks}$ = -0.03, 95% CI = [-0.06, -0.01], $pp_{>0}$ < .002). These results remained intact after accounting for choice difficulty. This was done by including the absolute subjective difference between the potential money and shock amounts (extracted from the best-fitting computational model, see main text **Results**) on each trial as an additional fixed effect(Krajbich, Bartling, Hare, & Fehr, 2015) (no random slope). Importantly, in this analysis we also observed a significant trial-by-trial relationship between response times and general heart rate deceleration (**Supplemental Figure S2A**; $B_{HR} = 0.03$, 95% CI = [0.01, 0.06], $pp_{<0} < .006$).

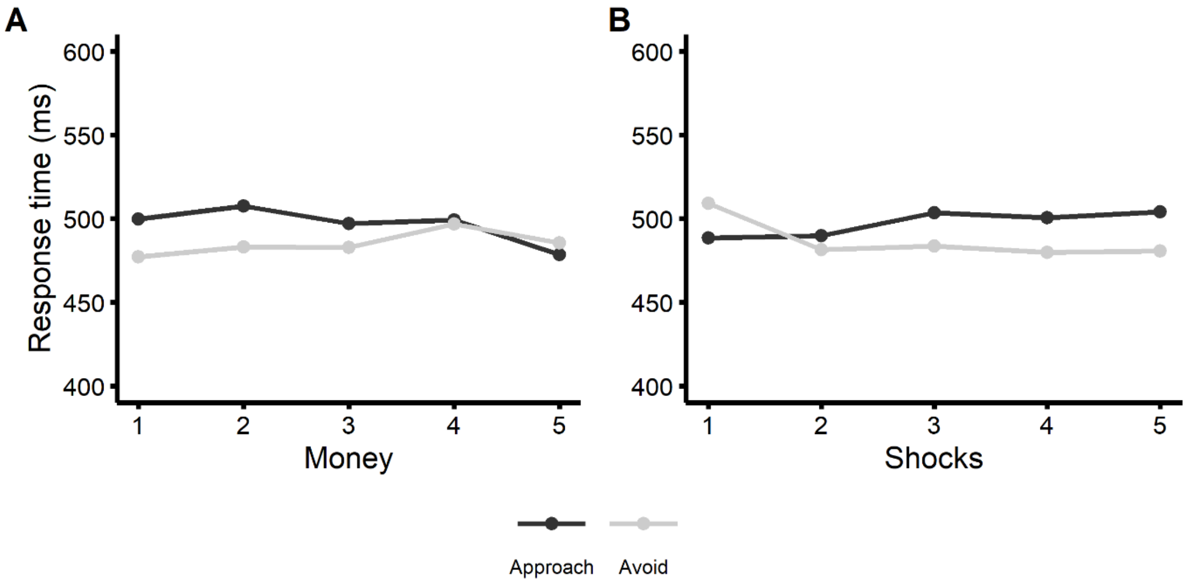


***Supplementary Figure S1. Task effects on response times.*** *Overall, average response times were marginally faster for avoid choices (**A, B**). Interestingly, increased money was associated with faster responses for approach choices (**A**), whereas an increase in shocks was associated with faster avoid choices (**B**).*

***Supplementary psychophysiological results***

As mentioned in the main text, the mixed-effects model on response times discussed above also revealed a significant trial-by-trial relationship between response times and general heart rate deceleration (**Supplemental Figure S2A**; $B_{HR} = 0.03$, 95% CI = [0.01, 0.06], $pp_{<0} < .007$).

To investigate the sensitivity of body sway to reward and punishment, we analyzed the body sway (BS) signal using a mixed-effects model with main and interaction effects of money and shock amounts. This showed a significant general decrease of BS in the time window of interest, relative to baseline (**Supplementary Figure S3A**; $B_{Intercept} = -0.25$, 95% CI = [-0.31, -0.19], $pp_{>0} < .001$). This reduction was however not moderated by money (95% CI = [-0.05, 0.03], $pp_{<0} = .688$) or shock amounts (95% CI = [-0.06, 0.01], $pp_{>0} = .096$).

Finally, trial-by-trial heart rate deceleration was significantly correlated to trial-by-trial body sway reduction **Supplementary Figure S2B**).

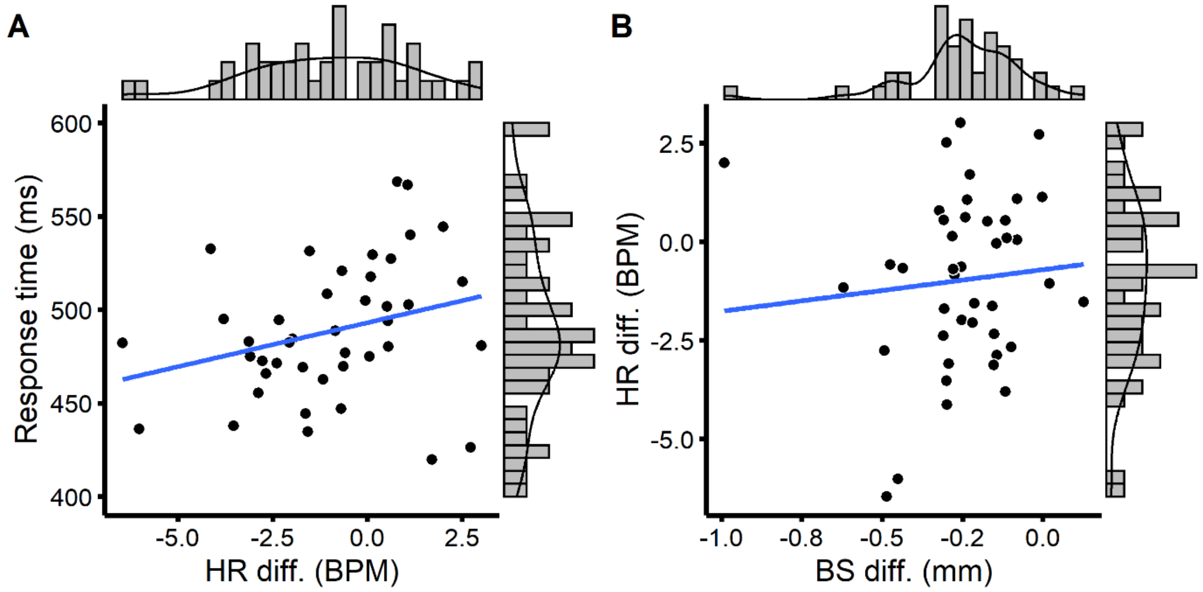


***Supplementary Figure S2. Physiological correlates.*** *General heart rate (HR) deceleration relative to pre-trial baseline (see main text Figure 4) was associated with faster response times (**A**). Moreover, trial-by-trial heart rate and body sway reductions were positively related (**B**). Solid vs. dashed regression lines reflect significant vs. non-significant relationships.*

**Supplementary computational modeling results**

***Robustness check – computational modeling of approach-avoidance choices with body sway as freeze index***

To check the robustness of the model comparison procedure, the same models as described in the main text were fitted again with body sway as alternative freeze index. This yielded the same winning model as when using heart rate as index (**Supplementary Figure S3B**).

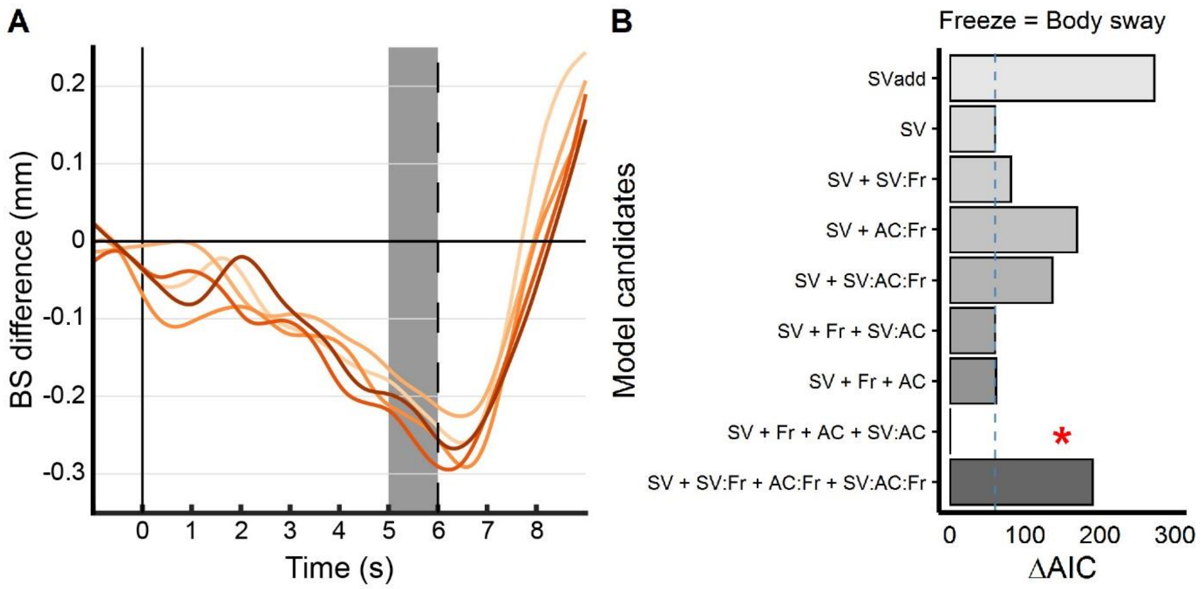


***Supplementary Figure S3. Postural freezing.*** *Body sway reduced significantly (i.e., postural freezing) during anticipation of approach-avoidance decisions (**A**). Using body sway as an index of freezing in our computational models yielded the similar results; the winning model was the same as when using heart rate (**B**).*

***Simulation of approach-avoidance choices***

We performed simulations of approach-avoidance choices (**Supplementary Figure S4**) to further understand the positive relationship between freeze-related bradycardia and the $\beta_{SV:AC}$ parameter (main text **Figure 3C**). To this end, we let the SV+AC+SV:AC model predict choices (approach/avoid) for all money, shock, and action context (passive/active) conditions while systematically varying the value of the $AC$ and $\beta_{SV:AC}$ parameters. More specifically, $AC$ was varied to simulate active ($AC$ = -0.8) and passive passive ($AC$ = 0.8) subjects, for both negative (-1) and positive (1) values of $\beta_{SV:SC}$, resulting in four separate simulations. For simplicity, all other parameters were set to have no influence on the choice (i.e., $\theta$ = 1, $\alpha_m$ = 1, $\alpha_s$ = 1, $\beta_{m:s}$ = 0).

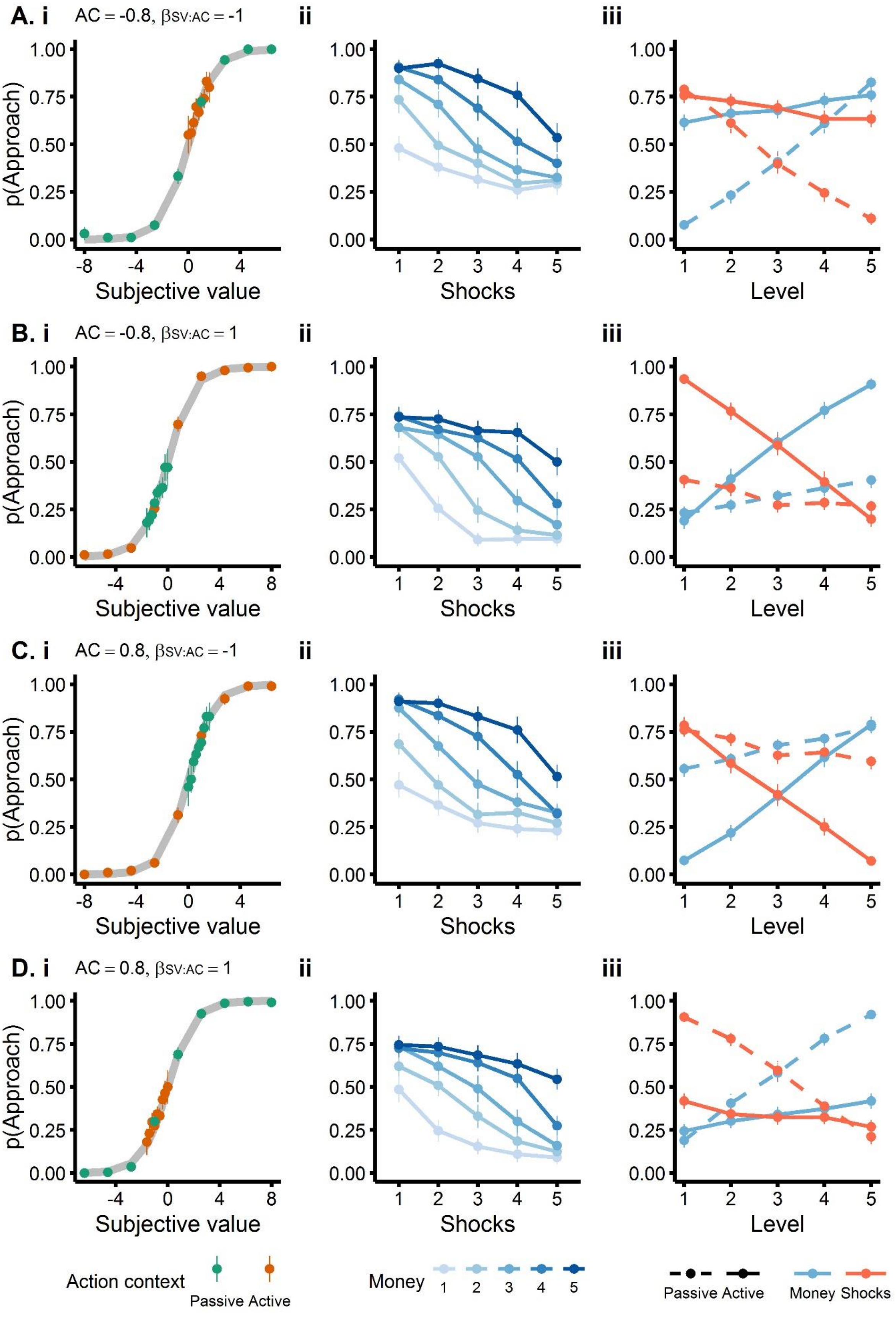
A. i
AC = -0.8, βSV:AC = -1
ii
iii
B. i
AC = -0.8, βSV:AC = 1
ii
iii
C. i
AC = 0.8, βSV:AC = -1
ii
iii
D. i
AC = 0.8, βSV:AC = 1
ii
iii
p(Approach)
Subjective value
Shocks
Level
Action context
Passive Active
Money
1 2 3 4 5
Passive Active
Money Shocks

***Supplementary Figure S4. Simulation of approach-avoidance choices.*** *Approach-avoidance choices were simulated for active (**A, B**) and passive tending (**C, D**) subjects, and for negative vs. positive subjective value by action context interactions (SV:AC). Panels in the left column (**i**) plot the predicted probability to approach as a function of the total subjective value per trial, with each dot representing the average choice of a single simulated individual plotted separately for active (orange) and passive action contexts (green). The middle and right columns (**ii, iii**) display the simulated proportion of approach choices as a function of varying money and shock amounts (1-5; **ii**) and action context (passive vs. active; **iii**). Error bars represent one standard error of the mean (SEM).*

***Inspection of fitted parameter estimates***

The fitted computational models produced no boundary fits, as is demonstrated in **Supplementary Table S1**.

**Supplementary Table S1**

*Summarized parameter estimates of best fitting model (SV+Fr+AC+SV:AC)*

| Parameter | Constraint | Median<br>HR \| BS | Interquartile range<br>HR \| BS | Min – Max<br>HR \| BS |
|---|---|---|---|---|
| $\alpha_m$ | -100 – 100 | 0.31 \| 0.32 | 0.28 \| 0.26 | -0.30 – 2.13 \| 0.001 – 2.10 |
| $\alpha_s$ | -100 – 100 | 0.39 \| 0.37 | 0.43 \| 0.42 | -0.16 – 1.53 \| -0.14 – 1.40 |
| $\beta_{m:s}$ | -100 – 100 | 0.10 \| 0.08 | 0.41 \| 0.40 | -0.93 – 2.55 \| -0.93 – 1.31 |
| $\beta_{Fr}$ | -100 – 100 | 0.003 \| -0.02 | 0.02 \| 0.09 | -0.04 – 0.07 \| -0.27 – 0.22 |
| AC | -100 – 100 | -0.37 \| -0.02 | 0.19 \| 0.18 | -2.05 – 0.30 \| -0.89 – 0.22 |
| $\beta_{SV:AC}$ | -100 – 100 | 0.40 \| -0.09 | 4.60 \| 5.53 | -11.55 – 80.84 \| -83.90 – 61.66 |

*Note.* Interquartile range is computed as the difference between the 3$^{rd}$ and 1$^{st}$ quantile. HR = heart rate, BS = body sway, m = money, s = shocks, Fr = umbrella term for freeze measures: either HR or BS, SV = subjective value, AC = action context.

**Supplementary control analyses**

***Manipulation check – effect of potential reward and threat on physiological arousal***

Although skin conductance was initially only measured for control analyses (see below), we performed a manipulation check to see whether our money and shock manipulations affected psychophysiological arousal. To that end, we ran a mixed-effects model on the skin conductance response (SCR) as a function of the potential money and shock amounts, and their interaction. To accommodate for the non-normal and zero-inflated shape of the SCR distribution, we used a hurdle model with a lognormal distribution to separately estimate the effect of our predictors on the probability of a non-zero response (i.e., the hurdle), and the amplitude of the SCR when it is non-zero.

This analysis revealed that both higher money and shock amounts led to increased probabilities of evoking a non-zero response ($B_{hu\_money}$ = -0.14, 95% CI = [-0.24, -0.05], $pp_{>0}$ < .004; $B_{hu\_shocks}$ = -0.27, 95% CI = [-0.38, -0.15], $pp_{>0}$ = .000) as well as increased skin conductance responses in general (**Supplementary Figure S5**; $B_{money}$ = 0.04, 95% CI = [0.01, 0.06], $pp_{<0}$ < .007; $B_{shocks}$ = 0.06, 95% CI = [0.03, 0.09], $pp_{<0}$ < .001). Our money and shock manipulations thus successfully invoked psychophysiological arousal.

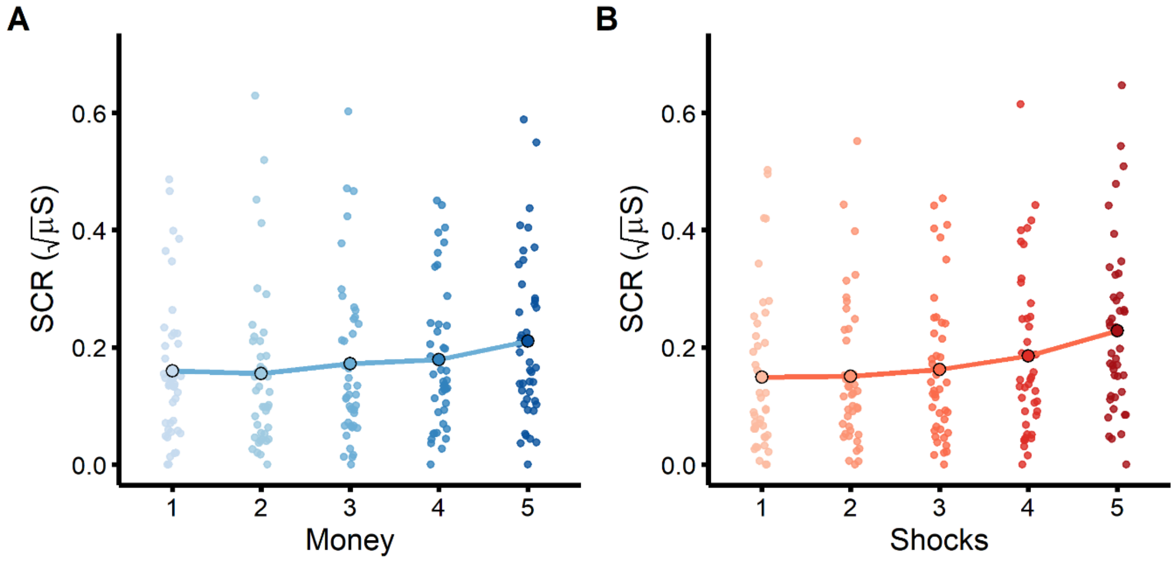


***Supplementary Figure S5. Effects of potential money and shock amounts on psychophysiological arousal.*** *Increased potential money (**A**) and shock (**B**) amounts are both associated with greater skin conductance responses (SCR). Lines and large dots represent overall means per money and shock amount; small dots represent individual subject means.*

***Control mixed-effects models – sympathetic influence and covariates***

To assess the robustness of our findings, we performed some control analyses. For the mixed-effects models, we first wanted to control for potential sympathetic influences on our dependent variables. To that end, we re-ran the three main mixed-effects models (on choice, response type, and response time) with skin conductance (SCR) as an additional fixed effect (no random slope). The results of these models were qualitatively identical to those reported in the main text. Then, as a general robustness check, we re-ran the same models after excluding two participants that showed very little variability in their choice data (i.e., <5 approach or avoid choices) and while controlling for trait anxiety and depression scores (Beck, Steer, & Brown, 1996; Spielberger, Gorsuch, Lushene, Vagg, & Jacobs, 1983) and gender (female/male). These models again yielded the same conclusions as reported above, except for one effect: the trend effect of choice on response time now became significant (i.e., faster response times for avoid compared to approach choices; $B_{choice}$ = -0.04, 95% CI = [-0.07 -0.002], $pp_{>0}$ = .02, note that critical $\alpha$ = .025). There were no effects of trait anxiety, depression, or gender.

For the computational models, we also controlled for potential sympathetic influence by running the same models with SCR as an extra predictor. This yielded the same conclusions with regards to the best fitting model.

Together, these control analyses confirm the robustness of the results reported in the main text.
